# Hemodynamic response function in resting brain: disambiguating neural events and autonomic effects

**DOI:** 10.1101/028514

**Authors:** Guo-Rong Wu, Daniele Marinazzo

**Affiliations:** Key Laboratory of Cognition and Personality, Faculty of Psychology, Southwest University, Chongqing, China.; Department of Data Analysis, Faculty of Psychology and Educational Sciences, Ghent University, Ghent, Belgium.

## Abstract

It has been shown that resting state brain dynamics can be characterized by looking at sparse blood-oxygen-level dependent (BOLD) events, which can be retrieved by point process analysis. Cardiac activity can also induce changes in the BOLD signal, thus affect both the number of these events and the mapping between neural events and BOLD signal, namely the hemodynamic response. To isolate neural activity and autonomic effects, we compare the resting state hemodynamic response retrieved by means of a point process analysis with and without deconvolving the cardiac fluctuations. Brainstem and the surrounding cortical area (such as precuneus, insula etc.) are found to be significantly affected by cardiac pulses. Methodological and physiological implications are then discussed.

## 1. Introduction

There is growing evidence indicating that discrete BOLD events govern the brain dynamic at rest (Deco and Jirsa 2012, Tagliazucchi, Balenzuela et al. 2012, Petridou, Gaudes et al. 2013, Wu, Liao et al. 2013). Relevant features of spontaneous neural activity could be therefore indirectly derived from these specific BOLD events. It has been shown that these events can be revealed by point processes analysis (PPA) (Tagliazucchi, Balenzuela et al. 2012). The core idea of PPA in this context is to isolate events in the BOLD time series (for example peaks in the standardized time series) and to look at their spatial and temporal distribution. Compared to static functional connectivity (FC) maps constructed from correlations between the whole time series, the FC maps derived by PPA appear to be similar but carry more information on the brain dynamics (Tagliazucchi, Balenzuela et al. 2012, Liu and Duyn 2013). Moreover, PPA is also used to retrieve the hemodynamic response function (HRF) at rest (Wu, Liao et al. 2013). Both FC and HRF can be employed to draw inferences on behavioral states and distinguish healthy and diseased populations (Handwerker, Gonzalez-Castillo et al. 2012, Barkhof, Haller et al. 2014). However, their statistical power is sensitive to non-neuronal artifacts. As the BOLD signal is a measurement of changes in blood flow, oxygenation, and volume (Ogawa, Lee et al. 1990), these changes may be caused by neuronal activity through neurovascular coupling, or arise from any other physiological process that affect blood oxygenation or volume (Birn 2012). Thus the precise neural underpinning of BOLD point processes is still not fully understood. Cardiac pulsations are among the most common physiological fluctuations that contribute to BOLD signal (Glover, Li et al. 2000, Shmueli, van Gelderen et al. 2007, de Munck, Goncalves et al. 2008, Chang, Cunningham et al. 2009). They contain both physiology-related spontaneous neuronal activity and non-neural fluctuations (Shmueli, van Gelderen et al. 2007), Birn (2012), (Thayer, Ahs et al. 2012, Garfinkel, Minati et al. 2014). Therefore it is critical to differentiate neural driven BOLD point process from confounding physiological BOLD point process of non-neuronal origin.

Spontaneous cardiac activity exhibits low frequency fluctuations and overlaps with frequency bands of interest in BOLD fluctuations (<0.1 Hz). Recent studies have shown that these nuisance confounds can significantly alter FC maps of the intrinsic brain networks, such as the default mode network (Shmueli, van Gelderen et al. 2007, Chang, Cunningham et al. 2009, Birn, Cornejo et al. 2014). Accordingly, it is expectable that the spatial and temporal distribution of point processes will also be affected. To date, it’s still not clear to what extent the cardiac pulsations affect the hemodynamic response retrieved by point process analysis, which is helpful for understanding the physiological foundation of functional coupling among brain regions (Valdes-Sosa, Roebroeck et al. 2011).

A number of methods have been developed to reduce cardiac variations induced fluctuations in the BOLD signal (Glover, Li et al. 2000, Shmueli, van Gelderen et al. 2007, Chang, Cunningham et al. 2009). Retrospective image space correction of physiological noise (RETROICOR) is one of the most employed methods to correct the cardiac pulsations (Glover, Li et al. 2000). However, it only filters cardiac cyclic effects aliased in the fMRI signal, while the cardiac-related low-frequency fluctuations remain in the data. The time-shifted heart rate (HR) time series was introduced to account for more variance in BOLD signal that the one induced by cardiac fluctuations (Shmueli, van Gelderen et al. 2007, Chang, Cunningham et al. 2009).

In the present study, to determine whether the spontaneous point processes hemodynamic response (HDR) reflects purely central processes (such as neural or astrocytic control), or if the HDR is affected by changes in cardiac activity (i.e. autonomic activity), resting state hemodynamic response function (HRF) is used to investigate the link between heart rate and brain activity at rest. The combination of HR and RETROICOR is employed to deconvolve the cardiac activity influence (Chang, Cunningham et al. 2009). Then spontaneous point process HRF maps are retrieved from the residual BOLD signal. Quantitative and test-retest analysis on HRF map with and without removing cardiac pulse is further performed.

## 2. Materials and Methods

### Data acquisition

The 7-Tesla resting-state (rs) fMRI test-retest dataset used in this study has been publicly released by the Consortium for Reliability and Reproducibility (CoRR) project (Gorgolewski, Mendes et al. 2015). Twenty-two participants (10 women) were scanned during two sessions spaced one week apart. Their age ranged from 21 to 30 with mean 25.1, one left handed subject was excluded, resulting in all right handed subjects). Each session includes two 1.5 mm isotropic whole-brain resting state scans (TR=3.0s, TE=17ms) and gradient echo field map. Structural images were acquired by 3D MP2RAGE sequence. Physiological data (respiratory and cardiac traces) was simultaneously recoded for each rs-fMRI scan.

### Data processing

All structural images in both datasets were manually reoriented to the anterior commissure and segmented into grey matter, white matter, and cerebrospinal fluid, using the standard segmentation option in SPM 12. Resting-state fMRI data preprocessing was subsequently carried out using both AFNI and SPM12 package. First, the EPI volumes of each run were corrected for the temporal difference in acquisition among different slices, and then the images were realigned to the first volume of the first run. The gradient echo field map was processed to create a voxel displacement map and used to correct the realigned images for geometric distortion. The resulting volumes were then despiked using AFNI’s 3dDespike algorithm to mitigate the impact of outliers. The mean BOLD image across all realigned volumes was coregistered with the structural image, and the resulting warps applied to all the despiked BOLD volumes. Finally all the coregistered BOLD images were spatially normalized into MNI space and smoothed (8 mm full-width half-maximum). The physiological data were down-sampled to 100 Hz.

### Cardiac fluctuation models

Two cardiac fluctuation models were constructed to account for components related to 1) cardiac phases (CP), 2) heart rate (HR). The motion parameters (MP, obtained in the realigning step) and respiration are also included to account for the physiological noise influences: 1) respiratory phases (RP) and the interaction effects between CP and RP (InterCRP), 2) respiratory volume (RV). Models for cardiac and respiratory phases and their interaction effects were based on RETROICOR (Glover, Li et al. 2000) and its extension (Harvey, Pattinson et al. 2008). Cardiac and respiratory response functions were employed to model heart rate and respiratory volume per time onto physiological process of the fMRI time series (Birn, Diamond et al. 2006, Shmueli, van Gelderen et al. 2007, Birn, Smith et al. 2008, Chang, Cunningham et al. 2009). For each subject, a set of 20 physiological regressors (i.e. 6 for CP, 8 for RP, 4 for InterCRP, RV, and HR) was created using the Matlab PhysIO toolbox for each EPI run. Cardiac fluctuation correction based on different combinations of these regressors was studied to investigate the effect of cardiac pulse, performing by a general linear model (GLM). The combinations are: 1) MP & RP & RV (MR-model), 2) MP & CP & RP & InterCRP & RV (MRC-model), 3) MP & HR & RV (MRH-model), 4) MP & CP & RP & InterCRP & HR & RV, i.e., all models (MRCH-model).

### Spontaneous point process event and HRF retrieval

We employ a blind deconvolution technique to retrieve spontaneous point process hemodynamic response function (HRF) from resting-state BOLD-fMRI signal (Wu, Liao et al. 2013). A linear time-invariant model for the observed resting state BOLD response is assumed (Wu, Liao et al. 2013). We hypothesize that a common HRF is shared across the various spontaneous point process events at a given voxel, resulting in a more robust estimation. After cardiac fluctuation correction, the BOLD signal at a particular voxel is given by:

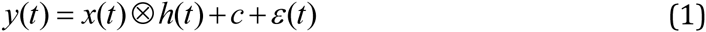

where *x*(*t*) is a sum of time-shifted delta functions centered at the onset of each spontaneous point process event and *h*(*t*) is the hemodynamic response to these events, *c* is a constant term indicating the baseline magnitude of the BOLD response, *ε*(*t*) represents additive noise and ⁫ denotes convolution. The noise errors are not independent in time due to aliased biorhythms and unmodelled neural activity, and are accounted for using an AR(p) model during the parameter estimation (we set p=2 in current study). Although no explicit external inputs exist in resting-state fMRI acquisitions, we still could retrieve the timing of these spontaneous events by means of the blind deconvolution technique (Wu, Liao et al. 2013). The lag between the peak of neural activation and the peak of BOLD response is presumed to be 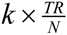 seconds (where 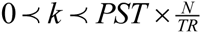 is the peristimulus time, PST). The timing set *S* of these resting-state BOLD transients is defined as the time points exceeding a given threshold around a local peak, is built in the following way: *S*{*i*} = *t_i_*, *y*(*t_i_*) ≥ *μ* & *y*(*t_i_*) ≻ *y*(*t_i_*–*τ*) & *y*(*t_i_*) ≻ *y*(*t_i_*–*τ*), where we set *τ* = 1,2 and *μ* = *σ* (i.e. the SD) in the current study. The exact time lag can be obtained by minimizing the mean squared error of equation (1), i.e. solving the optimization problem:

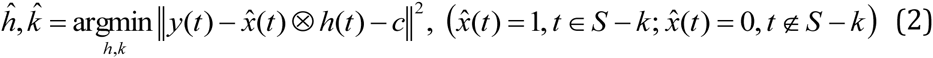

In order to avoid pseudo point process events induced by motion artifacts, a temporal mask with framewise displacement (FD)<0.3 was added to exclude these bad pseudo-event onsets from timing set *S* by means of data scrubbing (Power, Barnes et al. 2012). A smoothed finite impulse response (sFIR) model is employed to retrieve the spontaneous point process HRF shape (Goutte, Nielsen et al. 2000).

To characterize the shape of the hemodynamic response, three parameters, namely response height, time to peak, Full Width at Half Maximum (FWHM), were estimated, which could be interpretable in terms of potential measures for response magnitude, latency and duration of neuronal activity (Lindquist and Wager 2007).

After we retrieved the resting state HRF for each cardiac fluctuation correction model, the corresponding HRF parameters for each subject were individually entered into a random-effects analysis (one-way ANOVA within subjects, with three covariates (age, gender and mean FD) to identify regions which showed significant hemodynamic differences after cardiac fluctuation correction). Type I error due to multiple comparisons across voxels was controlled by familywise error rate (FWE).

### Test-retest reliability

The hemodynamic response has been shown to vary in timing, amplitude, and shape across brain regions and cognitive task paradigms (Miezin, Maccotta et al. 2000, Badillo, Vincent et al. 2013). Such variation is expected also for resting state. Previous studies have shown that physiological process account for significant variance, and enhance the test-retest reliability of functional connectivity map across subjects (Birn, Cornejo et al. 2014). In order to investigate the effect of cardiac fluctuation on the resting state HRF variability, a test-retest reliability analysis is further performed on the HRF parameters. As each subject was scanned in two sessions, each one with two runs, we could assess both inter- and intra-session reliability. Let BMS be the between-subject mean square and WMS the within-subject mean square. Then according to random effects model, an ICC value is defined by (Shrout and Fleiss 1979):

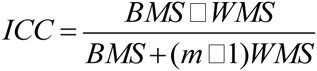

where *m* represents the number of repeated measurements of the voxelwise HRF parameter. ICCs were calculated for each voxel with intra-session scans and with inter-session scans, individually for each HRF parameter and cardiac fluctuation correction models. A prior functional parcellation of the cortex is applied to the ICC map in order to compute the mean ICC values and their standard deviations within sub-networks. This functional parcellation is composed of seven large-scale sub-networks: visual (VN), somatomotor (SMN), dorsal attention (DAN), ventral attention (VAN), limbic (LN), frontoparietal (FPN) and default network (DMN) (Yeo, Krienen et al. 2011).

## 3. Results

### Variance explained by different physiological components

Six head motion parameters and 20 physiological regressors are entered into mass univariate GLM analysis, and their efficacies are estimated by F test. Figure 1 shows the averaged fraction of variance explained by each regressors at voxelwise level over subjects. Most of high R-square values are distributed on the brainstem. For HR, the R-square values distribution is more homogeneous, and higher explained variance can also be found in cortical networks, such as SMN, VN and DMN. On average in brainstem, CP accounted for 7.9±5.3% of the variance, and InterCRP accounted for 4.02±2.67% of the variance, while HR only accounted for 1±0.5%.

**Figure 1.**
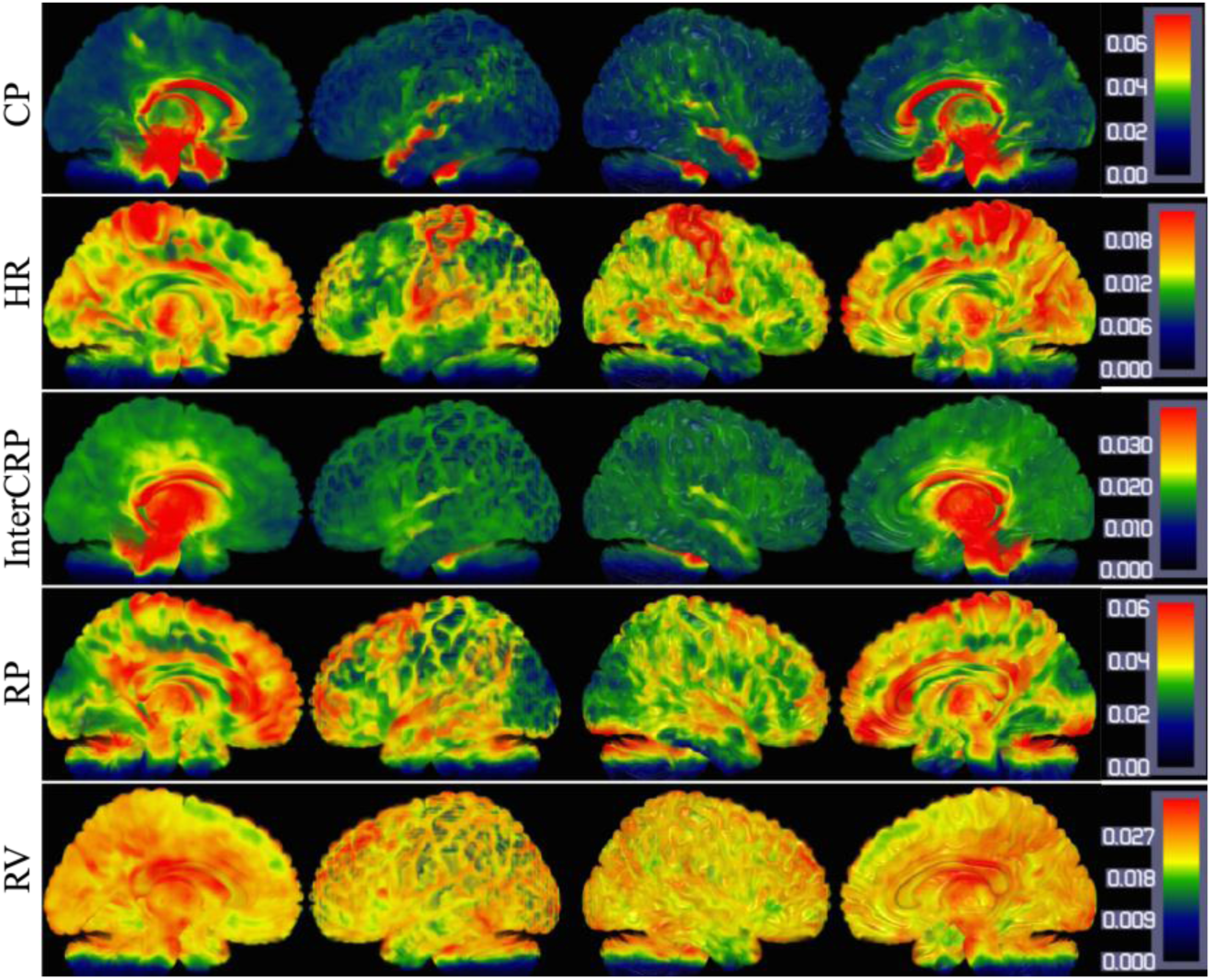
Spatial distribution of voxelwise R-squared values for different physiological components.

**Figure 2.**
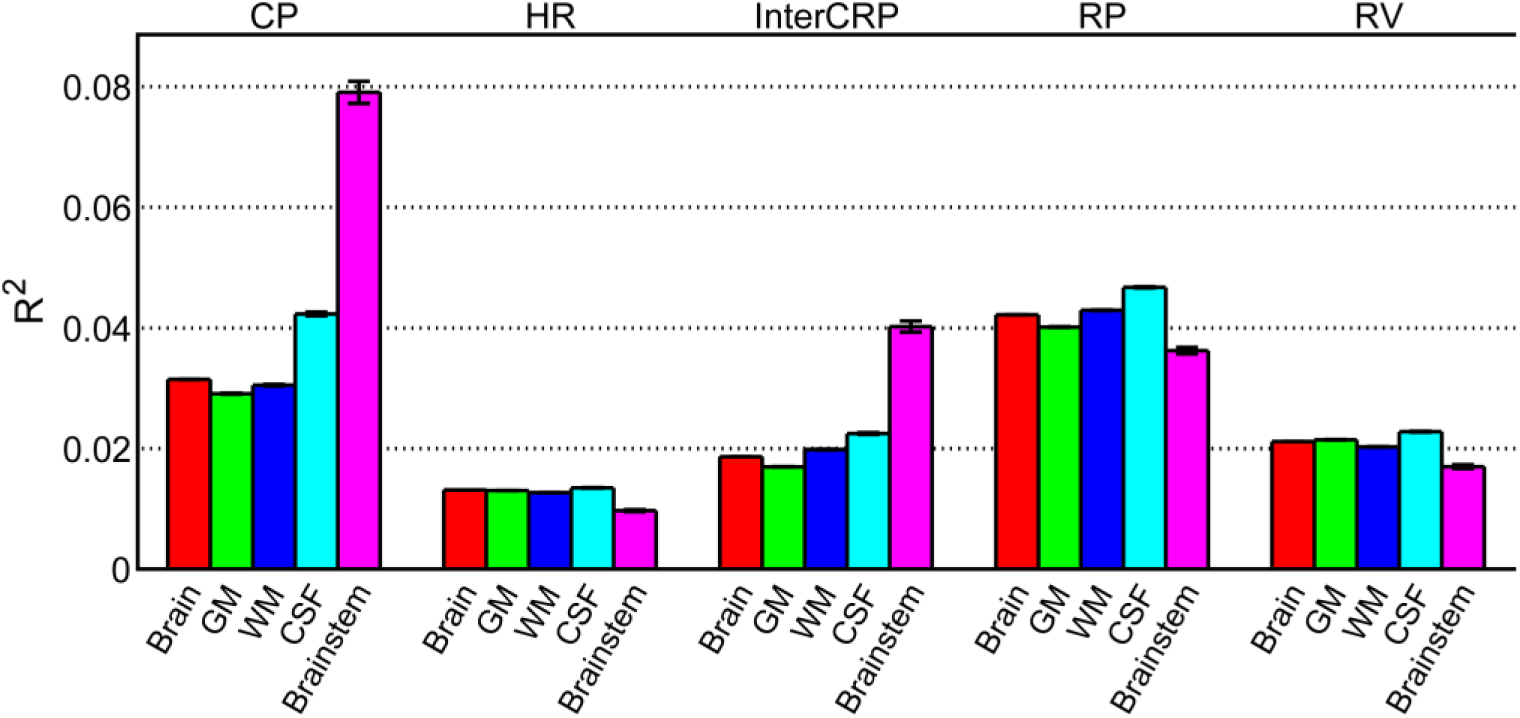
R-squared values for different physiological components, averaged over the entire brain, over all gray matter (GM), white matter (WM), cerebrospinal fluid (CSF), and brainstem. The black errorbar indicates the standard error.

### Spatial distributions of resting state HRF

HRF parameters of each voxel are estimated and mapped on the brain (Figure 3). The median maps of each HRF parameters exhibit spatial heterogeneity across different correction models (Figure 3). They present similar spatial distributions: higher response height is present in the occipital/frontal lobe and precuneus. As the temporal resolution is rather low (TR=3s) in this dataset, the median maps of FWHM and time to peak do not display consistent variations and are not shown here. The interested reader can find spatial maps of these parameters for shorter TRs in (Wu, Liao et al. 2013, Wu and Marinazzo 2015)

**Figure 3.**
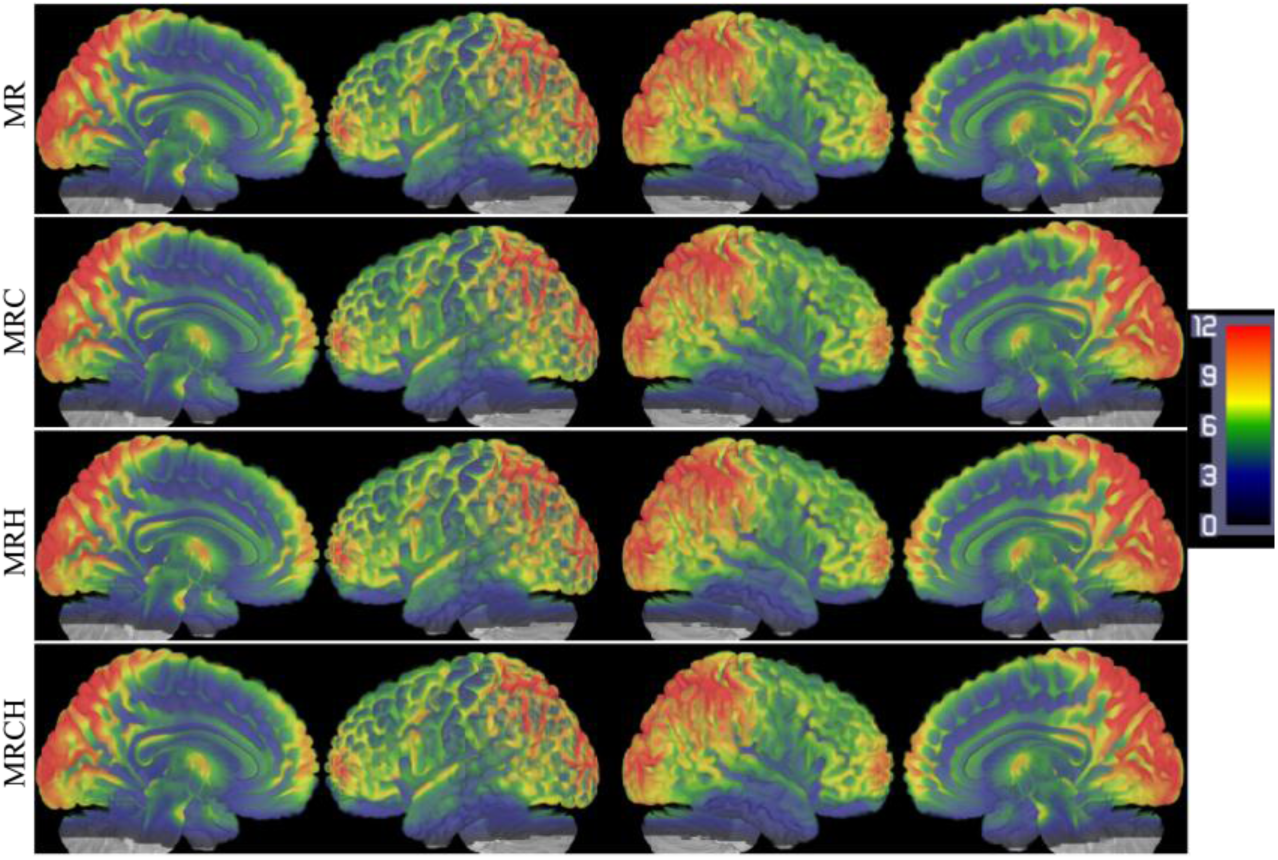
Median maps of HRF response heigth across subjects under different cardiac fluctuation correction model. The colorbar is same for all correction models.

### Group difference among the cardiac action models

Within-subjects ANOVA reveals that HRF response height is found to be significantly different across models. The main effect of the cardiac fluctuation model correction is mainly located in the brainstem and the surrounding pulsatile CSF regions and cortex: superior temporal gyrus, lingual gyrus, insula, parahippocampal gyrus, hippocampus, thalamus, putamen, caudate, amygdala, anterior/posterior cingulate, inferior/middle frontal gyrus, precuneus, and cuneus (Figure 4. p<0.05, FWE corrected). The HR gives a limited contribution to the variance of the point process HRF, while a significant contribution on brainstem’s variance comes from CP. This also appears evident looking at voxel-level HRFs shape averaged across subjects in brain stem and insula (Figure 5).

**Figure 4.**
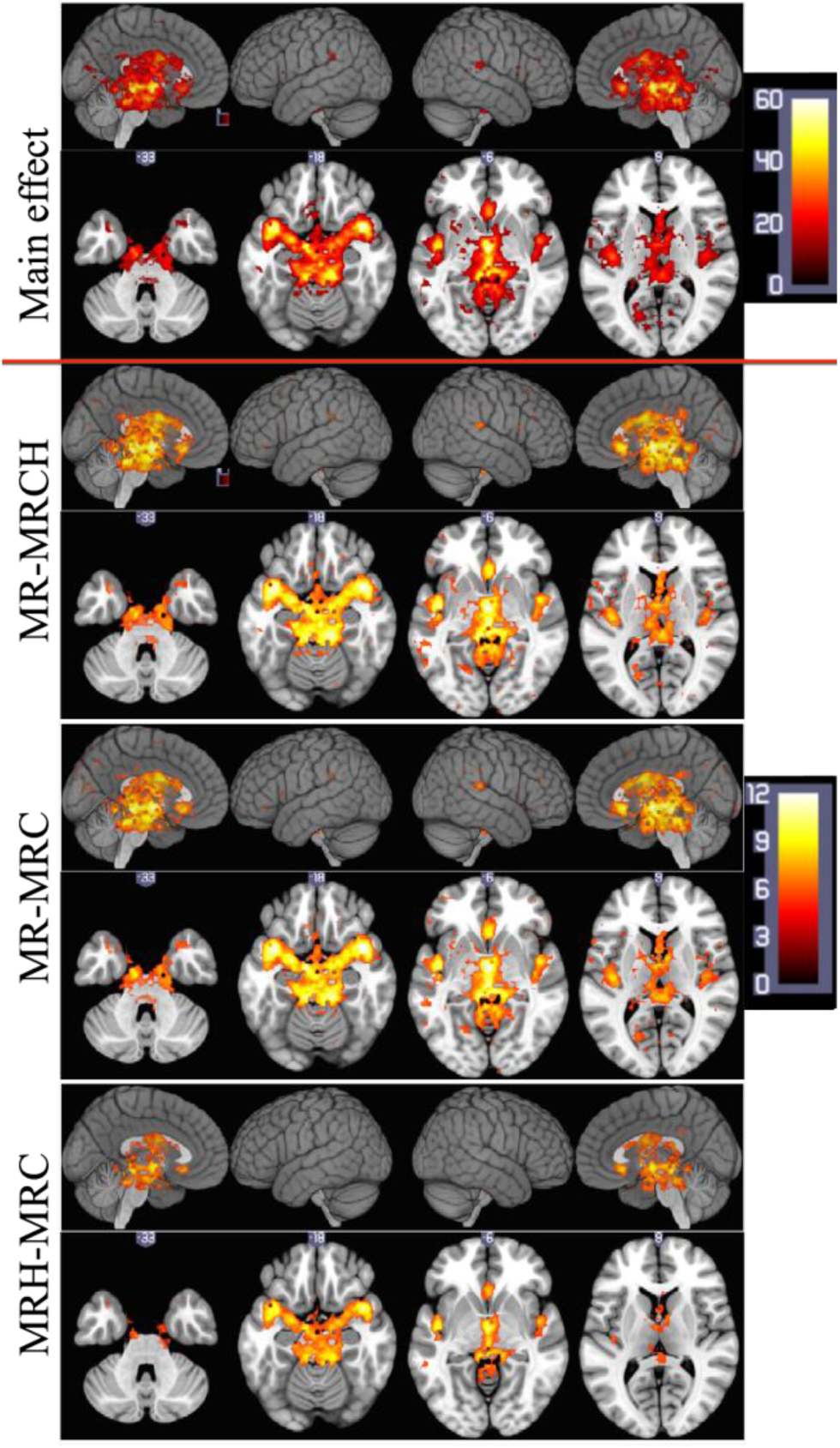
Top: main effect on HRF height among four cardiac fluctuation correction models. 2^nd^ row: differences in HRF response height between MR- and MRCH-model correction. 3^rd^ row: differences in response height between MR- and MRC-model correction. Bottom: differences in response height between MRH- and MRC-model corrections. All results are reported for a p-value <0.05, FWE corrected. The colorbar is the same from the three lowest panels.

**Figure 5.**
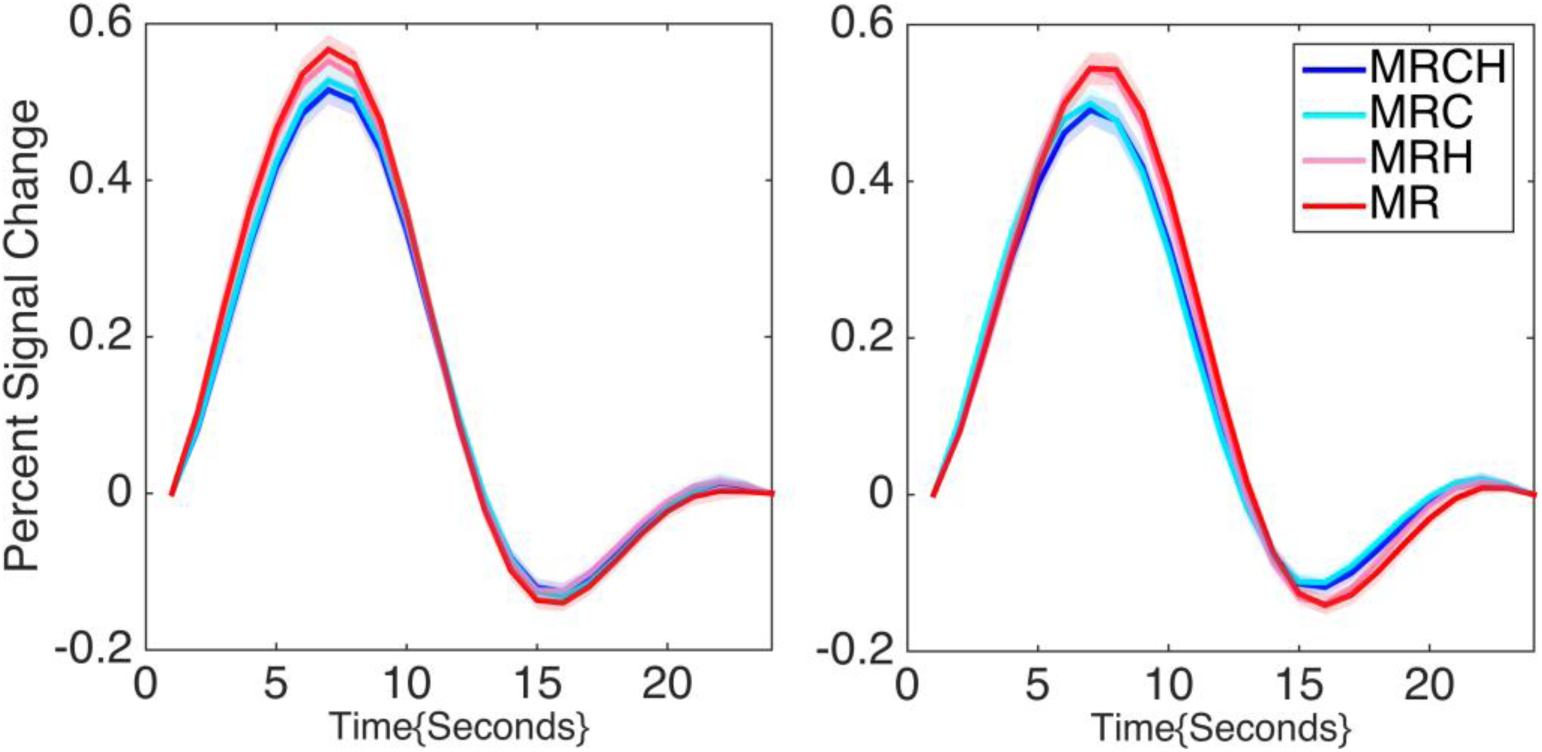
HRF at rest in the brainstem (right, MNI=[0, -27, -15]), and insula (left, MNI=[42, -9, 3]), retrieved from different cardiac fluctuation correction models. The colored shadow indicates the standard error.

### Test–retest reliability

As no difference was found in FWHM and time to peak in the ANOVA step, the test-retest analysis was only performed on the HRF response height. The effect of the corrections according to different cardiac fluctuation models are reported in Figure 6. There is no significant effect both within sessions and between sessions. According to the classifying criteria of ICC values (Sampat, Whitman et al. 2006), the intra-session of HRF response height shows excellent reliability (0.75~1), while inter-session mostly shows good reliability (0.6~0.75). Limbic network and brainstem show fair reliability (0.4~0.6). In contrast to intra-session, significant ICC reductions in inter-session are found in limbic, frontoparietal, default network, and brainstem (*p* < 0.05).

**Figure 6.**
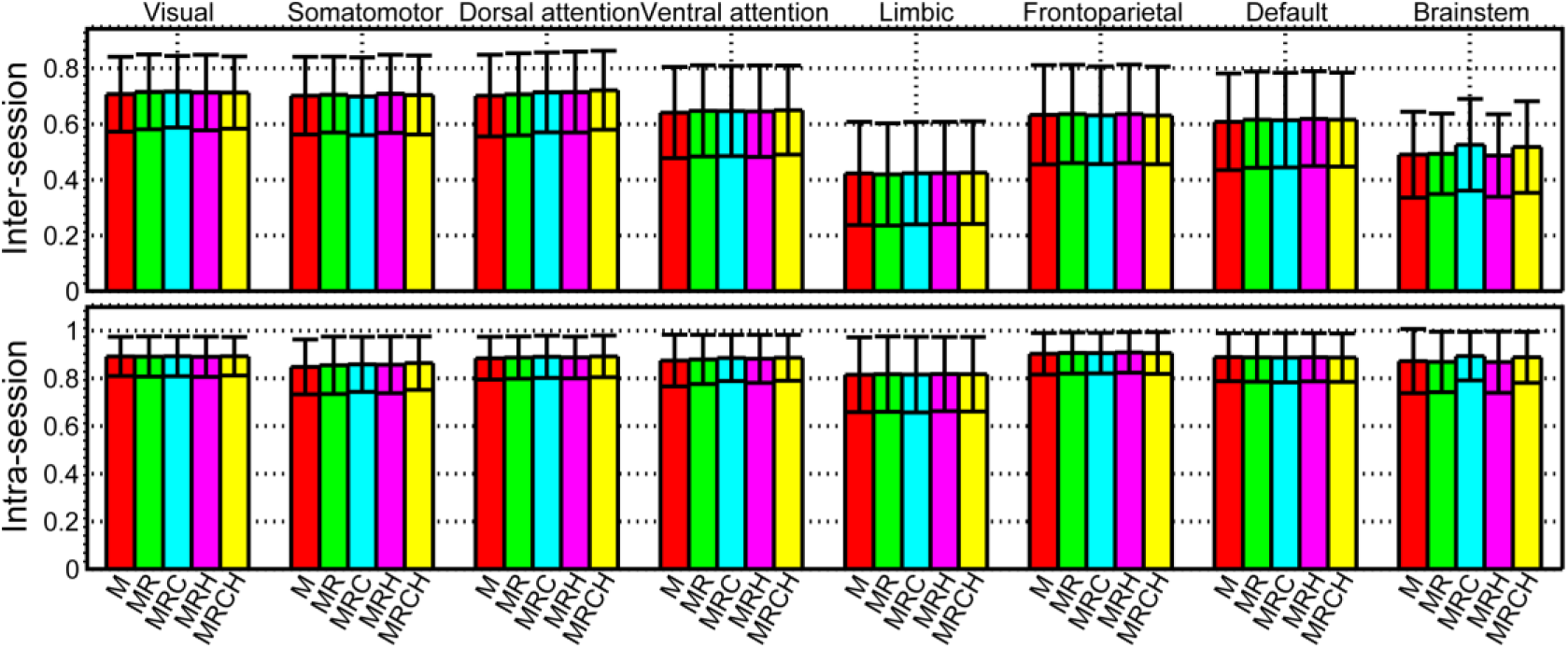
TRT reliability of HRF response height within seven subnetwork and brain stem. The black errorbars indicate the standard deviations.

## 4. Discussion

We investigated how cardiac fluctuations affect the resting state point process hemodynamic response. Brainstem and the surrounding cortical areas, comprising key regions of DMN (precuneus) and brain regions related to autonomic activity (such as insula and amygdala), are found to exhibit significant changes in hemodynamic response height. Specifically, cardiac phase accounts for the dominant variance alteration in point process HRF. A test-retest analysis revealed that cardiac fluctuations do not significantly change the reliability of point process HRF. Nonetheless, the higher inter-session variability of the height of point process HRF is mainly distributed on limbic network and brainstem, areas that are highly affected by cardiac activity. These results demonstrate that resting state point process HRF is a robust index and marker of brain activity, even without removing physiological fluctuations.

The neuroimaging evidence on brain-heart interactions mostly come from the regional cerebral blood flow, derived from PET or ASL (Restom, Behzadi et al. 2006, Thayer, Ahs et al. 2012), brain activity in PET/task fMRI or brain connectivity in resting state fMRI (Chang, Cunningham et al. 2009, Birn 2012, Thayer, Ahs et al. 2012). Our finding on spontaneous point process HRF related to cardiac fluctuations is consistent with these PET and fMRI studies. Our results show for the first time how HRF at rest is modulated by cardiac activity. Apart from brainstem, which is the most important integrative control center for autonomic nervous system function and plays an important role in the regulation of cardiac and respiratory function (Mendelowitz 1999, Beissner, Meissner et al. 2013), insular cortex is also posited to act as an integrator on the brain-heart axis (Nagai, Hoshide et al. 2010): it has a prominent role in limbic-autonomic integration and is involved in the perception of emotional significance (Augustine 1996); it also participates in visceral motor regulation, including blood pressure control, in cooperation with subcortical autonomic centers (Gianaros, Jennings et al. 2007, Napadow, Dhond et al. 2008, Lane, McRae et al. 2009). Other regions emerging from the current analysis are amygdala and anterior cingulate cortex (ACC), also involved in autonomic control (Critchley, Mathias et al. 2003, Chang, Metzger et al. 2013); the network consisting of insula, ACC, and amygdala has been described as crucial in the regulation of central autonomic nervous system (Critchley 2005). A human neuroimaging meta-analysis on electrodermal activity and high-frequency heart rate variability revealed that midbrain, insula and amygdala are associated with sympathetic and parasympathetic regulation; ACC and thalamus only correlated with sympathetic regulation, while dorsal posterior cingulated cortex, precuneus, superior temporal gyri and left temporal pole were associated to parasympathetic regulations (Beissner, Meissner et al. 2013).

The test-retest reliability is not significantly affected by all cardiac fluctuation models. A recent study reported a significantly decreased test-retest reliability in FC by physiological noise correction techniques (Birn, Cornejo et al. 2014). These results were explained by assuming that these physiological fluctuations are similar and reproducible within a subject across sessions, but to a lesser extent than between subjects. Another explanation given in the same study assumed that physiological noise correction could also remove the signal of interest. We found that HRF shape is only slightly modulated by cardiac pulsatility (Figure 5). This may explain that the spontaneous point process responses affected by cardiac fluctuation only account for small amounts of variance. On one hand, our result may confirm that these point processes are still preserved, i.e. the signal of interest is not removed. On the other hand, if the physiological fluctuations are similar and repeatable within a subject, no big HRF ICC differences after cardiac fluctuations correction is are expectable. Our results also indicate that HRF variability is much lower in lower-level perceptual networks and attention networks (DAN and VAN), and higher in emotional related network (LN), task-negative and task-positive networks (i.e. DMN and FPN). This may reflect intrinsic neural activity modulations in these networks after 1 week, and these neural point processes could also be associated with changes in autonomic activity (Fan, Xu et al. 2012).

Physiological fluctuations have been shown to be proportional to MRI signal strength (Kruger and Glover 2001): the physiological processes may therefore contribute much more to variance in BOLD signal with the current dataset, acquired at 7T. Apart from cardiac pulsations, respiration is another physiological fluctuation that has also been found to strongly modulate the resting state fMRI BOLD signal (Birn, Diamond et al. 2006, Birn, Smith et al. 2008). Respiration fluctuations will induce variations in arterial level of CO2, then cause either validations or vasoconstriction, resulting in blood flow and oxygenation changes (Van den Aardweg and Karemaker 2002). The fractions of variance explained by RV and HR on the resting state BOLD signal are found to mirror each other and to be partially co-localized, consistently with previous studies (Chang, Cunningham et al. 2009, Petridou, Schafer et al. 2009). As respiration and cardiac pulsations are tightly correlated (Pitzalis, Mastropasqua et al. 1997, Princi, Accardo et al. 2006), we first partial out respiration before investigating the impact or cardiac fluctuations on resting state point process based HRF. To reduce the computational cost and the bias in the linear estimation framework, we employ canonical functions for HR and RV hemodynamic response (Birn, Smith et al. 2008, Chang, Cunningham et al. 2009). The more flexible sFIR model could then minimize the risk of assumptions about the spontaneous point process HRF shape (Goutte, Nielsen et al. 2000). Moreover, sFIR model may also include the cardiac fluctuation related component in hemodynamic response, when it is not eliminated in BOLD signal. It’s well known that head motion is an unavoidable source of noise in the BOLD signal (Power, Mitra et al. 2014). To avoid motion-related artifacts contribution to point process detection, apart from adding MP as a nuisance regressor in GLM model, data scrubbing is performed (Power, Barnes et al. 2012), and Mean FD power of each subject was also included as a covariate for further statistical analysis (Van Dijk, Sabuncu et al. 2012). This procedure ensures that our findings are unlikely affected by motion artifact.

The precise estimation of HRF parameters, especially of FWHM and time to peak depends on the temporal resolution (Lindquist and Wager 2007). The quite low TR in current dataset could then have partially obscured findings related to the above mentioned parameters.

## Acknowledgments

This research was supported by the Natural Science Foundation of China (Grant No. 61403312), and the Fundamental Research Funds for the Central Universities (Grant No. 2362014xk04).

## References

Augustine, J. R. (1996). “Circuitry and functional aspects of the insular lobe in primates including humans.” Brain Res Brain Res Rev 22(3): 229–244.

Badillo, S., T. Vincent and P. Ciuciu (2013). “Group-level impacts of within- and between-subject hemodynamic variability in fMRI.” Neuroimage 82: 433–448.

Barkhof, F., S. Haller and S. A. Rombouts (2014). “Resting-state functional MR imaging: a new window to the brain.” Radiology 272(1): 29–49.

Beissner, F., K. Meissner, K. J. Bar and V. Napadow (2013). “The autonomic brain: an activation likelihood estimation meta-analysis for central processing of autonomic function.” J Neurosci 33(25): 10503–10511.

Birn, R. M. (2012). “The role of physiological noise in resting-state functional connectivity.” Neuroimage 62(2): 864–870.

Birn, R. M., M. D. Cornejo, E. K. Molloy, R. Patriat, T. B. Meier, G. R. Kirk, V. A. Nair, M. E. Meyerand and V. Prabhakaran (2014). “The influence of physiological noise correction on test-retest reliability of resting-state functional connectivity.” Brain Connect 4(7): 511–522.

Birn, R. M., J. B. Diamond, M. A. Smith and P. A. Bandettini (2006). “Separating respiratory-variation-related fluctuations from neuronal-activity-related fluctuations in fMRI.” Neuroimage 31(4): 1536–1548.

Birn, R. M., M. A. Smith, T. B. Jones and P. A. Bandettini (2008). “The respiration response function: the temporal dynamics of fMRI signal fluctuations related to changes in respiration.” Neuroimage 40(2): 644–654.

Chang, C., J. P. Cunningham and G. H. Glover (2009). “Influence of heart rate on the BOLD signal: the cardiac response function.” Neuroimage 44(3): 857–869.

Chang, C., C. D. Metzger, G. H. Glover, J. H. Duyn, H. J. Heinze and M. Walter (2013). “Association between heart rate variability and fluctuations in resting-state functional connectivity.” Neuroimage 68: 93–104.

Critchley, H. D. (2005). “Neural mechanisms of autonomic, affective, and cognitive integration.” J Comp Neurol 493(1): 154–166.

Critchley, H. D., C. J. Mathias, O. Josephs, J. O’Doherty, S. Zanini, B. K. Dewar, L. Cipolotti, T. Shallice and R. J. Dolan (2003). “Human cingulate cortex and autonomic control: converging neuroimaging and clinical evidence.” Brain 126(Pt 10): 2139–2152.

de Munck, J. C., S. I. Goncalves, T. J. Faes, J. P. Kuijer, P. J. Pouwels, R. M. Heethaar and F. H. Lopes da Silva (2008). “A study of the brain’s resting state based on alpha band power, heart rate and fMRI.” Neuroimage 42(1): 112–121.

Deco, G. and V. K. Jirsa (2012). “Ongoing cortical activity at rest: criticality, multistability, and ghost attractors.” J Neurosci 32(10): 3366–3375.

Fan, J., P. Xu, N. T. Van Dam, T. Eilam-Stock, X. Gu, Y. J. Luo and P. R. Hof (2012). “Spontaneous brain activity relates to autonomic arousal.” J Neurosci 32(33): 11176–11186.

Garfinkel, S. N., L. Minati, M. A. Gray, A. K. Seth, R. J. Dolan and H. D. Critchley (2014). “Fear from the heart: sensitivity to fear stimuli depends on individual heartbeats.” J Neurosci 34(19): 6573–6582.

Gianaros, P. J., J. R. Jennings, L. K. Sheu, S. W. Derbyshire and K. A. Matthews (2007). “Heightened functional neural activation to psychological stress covaries with exaggerated blood pressure reactivity.” Hypertension 49(1): 134–140.

Glover, G. H., T. Q. Li and D. Ress (2000). “Image-based method for retrospective correction of physiological motion effects in fMRI: RETROICOR.” Magn Reson Med 44(1): 162–167.

Gorgolewski, K. J., N. Mendes, D. Wilfling, E. Wladimirow, C. J. Gauthier, T. Bonnen, F. J. Ruby, R. Trampel, P. L. Bazin, R. Cozatl, J. Smallwood and D. S. Margulies (2015). “A high resolution 7-Tesla resting-state fMRI test-retest dataset with cognitive and physiological measures.” Sci Data 2: 140054.

Goutte, C., F. A. Nielsen and L. K. Hansen (2000). “Modeling the haemodynamic response in fMRI using smooth FIR filters.” IEEE Trans Med Imaging 19(12): 1188–1201.

Handwerker, D. A., J. Gonzalez-Castillo, M. D’Esposito and P. A. Bandettini (2012). “The continuing challenge of understanding and modeling hemodynamic variation in fMRI.” Neuroimage 62(2): 1017–1023.

Harvey, A. K., K. T. Pattinson, J. C. Brooks, S. D. Mayhew, M. Jenkinson and R. G. Wise (2008). “Brainstem functional magnetic resonance imaging: disentangling signal from physiological noise.” J Magn Reson Imaging 28(6): 1337–1344.

Kruger, G. and G. H. Glover (2001). “Physiological noise in oxygenation-sensitive magnetic resonance imaging.” Magn Reson Med 46(4): 631–637.

Lane, R. D., K. McRae, E. M. Reiman, K. Chen, G. L. Ahern and J. F. Thayer (2009). “Neural correlates of heart rate variability during emotion.” Neuroimage 44(1): 213–222.

Lindquist, M. A., and T. D. Wager (2007). “Validity and power in hemodynamic response modeling: a comparison study and a new approach.” Human brain mapping 28: 764–784.

Liu, X. and J. H. Duyn (2013). “Time-varying functional network information extracted from brief instances of spontaneous brain activity.” Proc Natl Acad Sci U S A 110(11): 4392–4397.

Mendelowitz, D. (1999). “Advances in Parasympathetic Control of Heart Rate and Cardiac Function.” News Physiol Sci 14: 155–161.

Miezin, F. M., L. Maccotta, J. M. Ollinger, S. E. Petersen and R. L. Buckner (2000). “Characterizing the hemodynamic response: effects of presentation rate, sampling procedure, and the possibility of ordering brain activity based on relative timing.” Neuroimage 11(6 Pt 1): 735–759.

Nagai, M., S. Hoshide and K. Kario (2010). “The insular cortex and cardiovascular system: a new insight into the brain-heart axis.” J Am Soc Hypertens 4(4): 174–182.

Napadow, V., R. Dhond, G. Conti, N. Makris, E. N. Brown and R. Barbieri (2008). “Brain correlates of autonomic modulation: combining heart rate variability with fMRI.” Neuroimage 42(1): 169–177.

Ogawa, S., T. M. Lee, A. R. Kay and D. W. Tank (1990). “Brain magnetic resonance imaging with contrast dependent on blood oxygenation.” Proc Natl Acad Sci U S A 87(24): 9868–9872.

Petridou, N., C. C. Gaudes, I. L. Dryden, S. T. Francis and P. A. Gowland (2013). “Periods of rest in fMRI contain individual spontaneous events which are related to slowly fluctuating spontaneous activity.” Hum Brain Mapp 34(6): 1319–1329.

Petridou, N., A. Schafer, P. Gowland and R. Bowtell (2009). “Phase vs. magnitude information in functional magnetic resonance imaging time series: toward understanding the noise.” Magn Reson Imaging 27(8): 1046–1057.

Pitzalis, M. V., F. Mastropasqua, F. Massari, A. Passantino, C. Forleo, G. Luzzi, P. Totaro, M. De Nicolo and P. Rizzon (1997). “Dependency of premature ventricular contractions on heart rate.” Am Heart J 133(2): 153–161.

Power, J. D., K. A. Barnes, A. Z. Snyder, B. L. Schlaggar and S. E. Petersen (2012). “Spurious but systematic correlations in functional connectivity MRI networks arise from subject motion.” Neuroimage 59(3): 2142–2154.

Power, J. D., A. Mitra, T. O. Laumann, A. Z. Snyder, B. L. Schlaggar and S. E. Petersen (2014). “Methods to detect, characterize, and remove motion artifact in resting state fMRI.” Neuroimage 84: 320–341.

Princi, T., A. Accardo and D. Peterec (2006). “Linear and non-linear assessment of heart rate variability in congenital central hypoventilation syndrome.” Biomed Sci Instrum 42: 434–439.

Restom, K., Y. Behzadi and T. T. Liu (2006). “Physiological noise reduction for arterial spin labeling functional MRI.” Neuroimage 31(3): 1104–1115.

Sampat, M. P., G. J. Whitman, T. W. Stephens, L. D. Broemeling, N. A. Heger, A. C. Bovik and M. K. Markey (2006). “The reliability of measuring physical characteristics of spiculated masses on mammography.” Br J Radiol 79 Spec No 2: S134–140.

Shmueli, K., P. van Gelderen, J. A. de Zwart, S. G. Horovitz, M. Fukunaga, J. M. Jansma and J. H. Duyn (2007). “Low-frequency fluctuations in the cardiac rate as a source of variance in the resting-state fMRI BOLD signal.” Neuroimage 38(2): 306–320.

Shrout, P. E., and J. L. Fleiss (1979). “Intraclass correlations: uses in assessing rater reliability.” Psychological bulletin 86(2): 420.

Tagliazucchi, E., P. Balenzuela, D. Fraiman and D. R. Chialvo (2012). “Criticality in large-scale brain FMRI dynamics unveiled by a novel point process analysis.” Front Physiol 3: 15.

Thayer, J. F., F. Ahs, M. Fredrikson, J. J. Sollers, 3rd and T. D. Wager (2012). “A meta-analysis of heart rate variability and neuroimaging studies: implications for heart rate variability as a marker of stress and health.” Neurosci Biobehav Rev 36(2): 747–756.

Valdes-Sosa, P. A., A. Roebroeck, J. Daunizeau and K. Friston (2011). “Effective connectivity: influence, causality and biophysical modeling.” Neuroimage 58(2): 339–361.

Van den Aardweg, J. G. and J. M. Karemaker (2002). “Influence of chemoreflexes on respiratory variability in healthy subjects.” Am J Respir Crit Care Med 165(8): 1041–1047.

Van Dijk, K. R., M. R. Sabuncu and R. L. Buckner (2012). “The influence of head motion on intrinsic functional connectivity MRI.” Neuroimage 59(1): 431–438.

Wu, G.-R., and D. Marinazzo (2015). Retrieving the Hemodynamic Response Function in resting state fMRI: methodology and applications, PeerJ PrePrints.

Wu, G. R., W. Liao, S. Stramaglia, J. R. Ding, H. Chen and D. Marinazzo (2013). “A blind deconvolution approach to recover effective connectivity brain networks from resting state fMRI data.” Med Image Anal 17(3): 365–374.

Yeo, B. T., F. M. Krienen, J. Sepulcre, M. R. Sabuncu, D. Lashkari, M. Hollinshead, J. L. Roffman, J. W. Smoller, L. Zollei, J. R. Polimeni, B. Fischl, H. Liu and R. L. Buckner (2011). “The organization of the human cerebral cortex estimated by intrinsic functional connectivity.” J Neurophysiol 106(3): 1125–1165.

